# Training-dependent gradients of timescales of neural dynamics in the primate prefrontal cortex and their contributions to working memory

**DOI:** 10.1101/2023.09.01.555857

**Authors:** Ethan Trepka, Mehran Spitmaan, Xue-Lian Qi, Christos Constantinidis, Alireza Soltani

## Abstract

Cortical neurons exhibit multiple timescales related to dynamics of spontaneous fluctuations (intrinsic timescales) and response to task events (seasonal timescales) in addition to selectivity to task-relevant signals. These timescales increase systematically across the cortical hierarchy, e.g., from parietal to prefrontal and cingulate cortex, pointing to their role in cortical computations. It is currently unknown whether these timescales depend on training in a specific task and/or are an inherent property of neurons, and whether more fine-grained hierarchies of timescales exist within specific cortical regions. To address these questions, we analyzed single-cell recordings within five subregions of the prefrontal cortex (PFC) of male macaques before and after training on a working-memory task. We found fine-grained but opposite gradients of intrinsic and seasonal timescales that mainly appeared after training. Intrinsic timescales decreased whereas seasonal timescales increased from posterior to anterior subregions within both dorsal and ventral PFC. Moreover, training was accompanied by increases in proportions of neurons that exhibited intrinsic and seasonal timescales. These effects were comparable to the emergence of response selectivity due to training. Finally, task selectivity accompanied opposite neural dynamics such that neurons with task-relevant selectivity exhibited longer intrinsic and shorter seasonal timescales. Notably, neurons with longer intrinsic and shorter seasonal timescales exhibited superior population-level coding, but these advantages extended to the delay period mainly after training. Together, our results provide evidence for plastic, fine-grained gradients of timescales within PFC that can influence both single-cell and population coding, pointing to the importance of these timescales in understanding cognition.

**Significance statement:** Recent studies have demonstrated that neural responses exhibit dynamics with different timescales that follow a certain order or hierarchy across cortical areas. While the hierarchy of timescales is consistent across different tasks, it is unknown if these timescales emerge only after training or if they represent inherent properties of neurons. To answer this question, we estimated multiple timescales in neural response across five subregions of the monkeys’ lateral prefrontal cortex before and after training on a working-memory task. Our results provide evidence for fine-grained gradients related to certain neural dynamics. Moreover, we show that these timescales depend on and can be modulated by training in a cognitive task, and contribute to encoding of task-relevant information at single-cell and population levels.

## Introduction

Considerable variability is present in neural responses to stimuli, rewards, and cognitive operations within all brain areas (Chaudhuri et al., 2015). Despite this heterogeneity, the dynamics of spontaneous fluctuations in cortical neural response exhibit a hierarchy of timescales (intrinsic timescales) that broadly corresponds to the cerebral cortex’s processing hierarchy (Honey et al., 2012; Goris et al., 2014; Murray et al., 2014; Song et al., 2023).

Specifically, neurons in sensory areas sensitive to tracking dynamic stimuli exhibit shorter timescales than neurons in the association cortex involved, for example, with maintaining information in working memory. This hierarchy of intrinsic timescales across cortex is accompanied by parallel hierarchies of timescales related to response to task events (seasonal timescales) and the memory of task-relevant signals such as previous choice and reward outcomes (Bernacchia et al., 2011; Murray et al., 2014; Spitmaan et al., 2020; Song et al., 2023). Interestingly, although the responses of individual cortical neurons exhibit multiple such timescales simultaneously, these timescales are not correlated across individual neurons within a single brain area, suggesting that they may have different roles in cortical computations (Spitmaan et al., 2020).

In addition, these neuronal timescales may reflect behavioral timescales related to integration of information over time according to the demands of a given cognitive task. For example, intrinsic timescales of cortical neurons have been shown to correlate with timescales of evidence accumulation in a perceptual decision-making task (Pinto et al., 2022) and have been hypothesized to play a key role in processing sensory input at different timescales (Golesorkhi et al., 2021). Moreover, intrinsic timescales seem to be predictive of information encoding in parietal and prefrontal neurons during a working-memory task with an overall better decoding performance for neurons with longer intrinsic timescales (Wasmuht et al. 2018; Cavanagh et al. 2018). On the other hand, timescales of neural activity tied to the memory of choice and reward outcomes (choice- and reward-memory timescales) have been shown to correlate with behavioral timescales of choice and reward integration in a value-based decision making task (Spitmaan et al., 2020).

How these timescales, which have been observed experimentally in trained animals during the execution of different cognitive tasks, appear in the first place is currently unknown. The concept of an “intrinsic” timescale implies that the timescale arises from biophysical properties that are inherent to a neuron (and its circuit) and thus are immutable, however, this idea has not been tested experimentally. On the other hand, timescales tied to task events (e.g., seasonal timescales) may have either emerged after task acquisition or may have relied on a preexisting timescale architecture as some modeling studies have suggested (Kim and Sejnowski, 2021).

Here, to address the above questions and test for possible fine-grained gradients of timescales related to different neural response dynamics, we estimated multiple timescales in responses of individual neurons in the prefrontal cortex (PFC) of monkeys recorded before and after training on a working-memory task. We focused on five subregions of the lateral PFC, anterior-dorsal, mid-dorsal, posterior-dorsal, anterior-ventral, and posterior-ventral, because these subregions were shown to exhibit distinct functional properties and capacity for plasticity (Riley et al., 2017, 2018). Overall, our results provide evidence for fine-grained gradients of intrinsic and seasonal timescales along the anterior-posterior axis. These gradients were further shaped by training in the task and were predictive of selectivity to task-relevant signals. However, the direction of gradients and their relationship to selectivity were opposite for the intrinsic and seasonal timescales with direct effects on population coding. More specifically, neurons with task-relevant selectivity exhibited longer intrinsic and shorter seasonal timescales that accompanied superior population-level encoding of cue stimulus. Notably, these advantages extended to the delay period between cue and sample presentations mainly after training, suggesting that such neurons could be selectively recruited for performing the working-memory task.

## Materials and Methods

### Experimental design and statistical analysis

The data analyzed in this study were previously described in Riley et al. (2018). Briefly, neuronal recordings were carried out in PFC while monkeys performed a passive-viewing task (during pre-training phase) and a spatial match/non-match working memory task (during post-training phase). The monkeys did not have any task experience prior to the experiment. Six male rhesus monkeys (ages 5–9 years) performed the pre-training task, and four of the six monkeys performed the post-training task.

In the pre-training task, a central fixation point was displayed for 1 s, followed by a cue stimulus for 0.5 s, a fixation point for 1.5 s (first delay period which we refer to as cue delay), a sample stimulus for 0.5 s, and a fixation point for another 1.5 s (second delay period). The first visual stimulus (cue stimulus) could appear at one out of nine possible locations around the fixation point (**Fig. 1b**), but the second visual stimulus (sample stimulus) was shown either at the same location as the cue or diametrically opposite the cue. Monkeys received a reward (fruit juice) if they maintained fixation during the trial, so the locations of stimuli had no behavioral relevance to the monkeys during the task.

**Figure 1.**
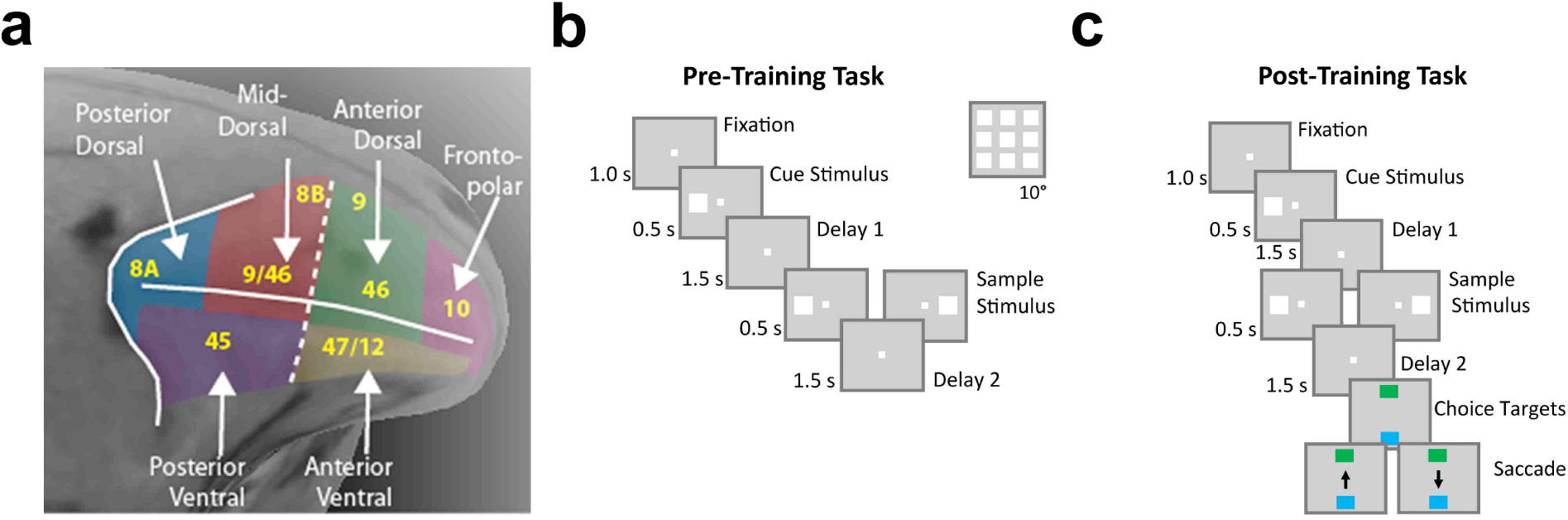
Schematic of the behavioral task and subregions in the lateral prefrontal cortex where neurons were recorded before and after training. **(a)** Subregions of the monkey lateral prefrontal cortex as defined in this study. **(b–c)** Timeline of a pre-training **(b)** and post-training **(c)** trial. At the onset of both trials, monkeys fixated on a central point and then were presented with a cue stimulus followed by a delay period (delay 1 = 1.5 sec). After the delay period, monkeys were presented with a sample stimulus followed by a second delay period (delay 2 = 1.5 sec). Cue and sample stimuli could appear at the nine locations surrounding the central point (inset in b). In pre-training trials, monkeys then received a juice reward for maintaining fixation on the central point throughout the trial **(b)**. In post-training trials, monkeys then made a saccade to either a green or blue square (or an H or diamond stimulus) to indicate that the cue and sample locations matched or did not match, respectively **(c)**.

The post-training task was similar to the pre-training task but required the monkeys to remember and compare the locations of cue and sample stimuli and provide a response. More specifically, after the second delay period, monkeys made a saccade to one of two target stimuli to report whether the locations of the cue and sample stimuli matched or did not match. Target stimuli were a green or blue square for two of the monkeys and an “H” or diamond for the other two monkeys. In contrast to the pre-training task, the monkeys received a juice reward only if they correctly indicated that the cue and sample stimuli matched or did not match.

Extracellular recordings were performed in areas 8, 9, 9/46, 45, 46, and 47/12 of the PFC while the monkeys performed the pre- and post-training tasks. Prior to the first recording, the monkeys underwent a magnetic resonance imaging scan so that electrode penetrations could be mapped onto PFC subregions. Recordings were performed using arrays of up to 8 glass- or epoxylite-coated tungsten electrodes (Alpha-Omega Engineering, Nazareth, Israel), and neurons were sampled in an unbiased fashion such that all units isolated from recordings were included in analyses.

During the experiment, eye position was monitored using an infra-red eye tracking system (model RK-716; ISCAN, Burlington, MA), and presentation of stimuli and data collection was handled with custom software implemented in MATLAB (Mathworks, Natick, MA) and PsychToolbox (Brainard, 1997). All experimental procedures followed guidelines by the U.S. Public Health Service Policy on Humane Care and Use of Laboratory Animals and the National Research Council’s Guide for the Care and Use of Laboratory Animals. All experimental procedures were reviewed and approved by the Wake Forest University Institutional Animal Care and Use Committee. For more details about the task design, neuronal recordings, and anatomical localization, see Riley et al. (2018).

Standard inferential statistics were used to identify changes in timescales in prefrontal subregions before and after training and to assess the relationship between timescales and selectivity. Statistical analyses of gradients of timescales along the anterior-posterior axis of PFC can be found in the **Results** describing **Figs. 3–4**. Two-sided Wilcoxon rank sum tests were used to test for changes in median intrinsic and seasonal timescales in prefrontal subregions before and after training (**Fig. 5**) and in neurons with and without task-relevant components (**Fig. 6**). Chi-squared tests were used to identify changes in the proportion of neurons that included a particular model component before and after training (**Fig. 5**). Spearman’s rank correlations were used to test for associations between timescales and variance explained by task-relevant components (**Fig. 7**).

Finally, details of the decoding analyses can be found in the preceding section and the **Results** describing **Figs. 8–9**.

### Estimating multiple timescales of neural response

We first briefly review the method for estimating multiple timescales via predicting spike counts developed by Spitmaan and colleagues (Spitmaan et al., 2020), which has been further modified and extended here. The goal of this method is to fit the spike count of individual neurons over multiple trials of an experimental session. The spike count is inherently a non-stationary time series because of the dependence of neural activity on task events and preceding neural activity on different timescales. One common method to predict non-stationary time series is a seasonal autoregressive with exogenous inputs model referred to as ARX that incorporates both an autoregressive component to capture the dependence of the outcome variable on its preceding values as well as exogenous components to capture the periodic effects of external events on the outcome variable (Seo and Lee, 2007; Seo et al., 2009; Box et al., 2015). To predict neural response in a given 50-msec time bin, Spitmaan et al. (2020) utilized a 2D-ARX model with two autoregressive components to capture the influence of activity in earlier epochs of the same trial (intrinsic component) and activity from the same epoch of previous trials (seasonal component), two exponential memory traces to capture the influence of choice and reward from previous trials, and two exogenous components to capture the influence of choice and reward in the current trial.

We modified and extended this model here by incorporating four exogenous components that capture the influence of the cue stimulus location, sample stimulus location, match vs non-match (match status), and the interaction between sample location and match status. We did not include memory traces for cue or sample location in the previous trials because our additional analyses suggested that the cue and sample stimulus location in past trials did not explain a substantial amount of variance in neural response. Similar to Spitmaan et al. (2020), we incorporated an intrinsic autoregressive component (*AR_intrinsic_*) that uses a weighted average of spike counts in the preceding *F* 50-msec time bins to predict spike count in the current 50-msec time bin as well as a seasonal autoregressive component (*AR_seasonal_*) that uses the spike count in the same time bin of *G* previous trials to predict spike count in the current time bin of the current trial. The coefficients of each *AR_intrinsic_* and *AR_seasonal_* component are then transformed to *τ_intrinsic_* and *τ_seasonal_*.

More formally, the model predicts the mean-subtracted firing rate in bin *n* of trial *k*, *y∼(n, k)* = *y(n, k) − E[y(n, k)]_k_*, using the following equation:

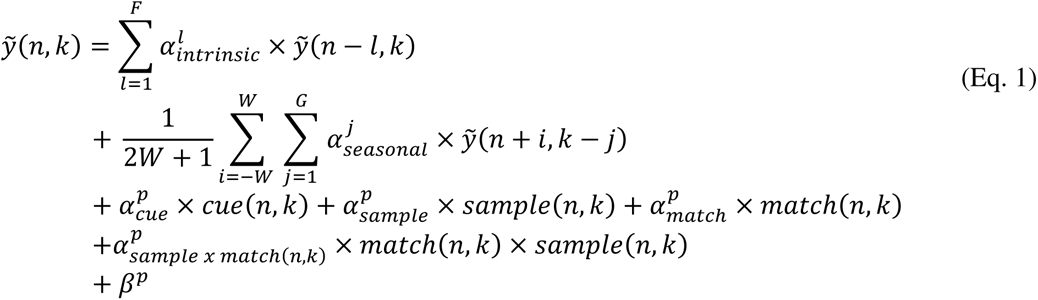

where 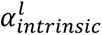 and 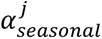 denote the autoregressive coefficients for intrinsic and seasonal fluctuations, respectively. The intrinsic autoregressive component is order *F*, and the seasonal autoregressive component is order *G*. The firing rate in previous trials used for the seasonal predictor is smoothed with a 250-msec smoothing window (i.e., *W = 2)* to reduce noise in the signal while preserving temporal specificity. 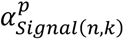 is the coefficient for the exogenous signal *signal(n, k)* during the current task period *p ∈*{fixation, cue stim., delay 1, sample stim., delay 2}. The cue exogenous signal is only present during the cue stimulus and delay 1 (cue delay) periods, and the sample, match, and interaction exogenous signals are only present during the sample stimulus and delay 2 periods. Five bias terms (*f]^p^)* are included in the model, one for each of the 5 task periods.

To determine the order of the intrinsic and seasonal autoregressive components, we computed the partial autocorrelation function (PACF) for intrinsic and seasonal time series and identified the time lag when the mean intrinsic and seasonal PACFs crossed the 95% confidence interval (**Fig. 2-1**) (Box et al. 2015). This results in order ten for the intrinsic component (*F* = 10) and order one for the seasonal component (*G* = 1).

The cue (sample) exogenous signal is a categorical variable with 9 categories corresponding to the 9 possible cue locations. Thus, 8 predictors for the cue (sample) exogenous signal were included in the model for each task period that the cue signal was present. The match signal is a categorical variable with 2 categories corresponding to match and non-match. Thus, one predictor for the match exogenous signal was included in the model for each task period that the match signal was present. 7 predictors for the sample x match signal were included in the model for each task period that the signal was present because a sample signal in the center location could only correspond to a match trial. The match signal and sample x match signal capture all forms of mixed selectivity (Rigotti et al., 2013; Dang et al. 2021) that could be present in the task because the sample stimulus was shown either at the same location as the cue or diametrically opposite the cue.

To detect multicollinearity among predictors in the model, we computed the Pearson correlations between all pairs of predictors for each neuron, then averaged the resulting correlation matrix across all pre-training neurons or all post-training neurons. We did not observe large correlations among predictors in the revised model before training or after training (mean r < 0.15) for all pairs of predictors with the exception of the sample, match, and sample x match predictors.

Correlations between sample, match, and sample x match predictors were expected because these predictors are non-zero in the same time intervals within a trial, and the sample x match predictors are derived from the sample and match predictors.

Above describes the full model for predicting spike counts with all components. To measure the contribution of each component to the full model, we also constructed more specific models by turning off a single autoregressive or exogenous component and refitting the model with all but one predictor.

### Model fitting procedure

Model parameters for the autoregressive components were determined by fitting the full model for each neuron using ordinary least-squares regression. To assess the predictive power of the full model, we fit the model using ten-fold cross-validated regression.

Specifically, we generated each instance of training data by randomly sampling 90% of all data (bins) for each neuron and then calculated fitting performance based on R-squared in the remaining 10% of data (test data). To determine the lower-bound on the contribution of each component to the neuron’s response, we fit the full model with a single component removed using ten-fold cross-validated regression. To determine the upper-bound on the contribution of each component to the neuron’s response, we fit models composed of only a single component using ten-fold cross-validated regression. All figures are based on parameter estimates from the full model unless otherwise specified. Models were fit to neural data in five seconds of each trial, from 1 second before cue presentation to the end of the second delay period. Moreover, models were fit only to data from completed trials in which the animal maintained fixation for the duration of the trial and made a saccade to a choice target (only in the post-training task). In the post-training task, both trials in which the animal made a correct or incorrect choice were included to maximize the number of sequential trials for estimation of seasonal timescales.

### Model selection procedure

A neuron includes a particular component if the addition of that component to the model results in a statistically significant increase in explained variance. More specifically, we computed the R^2^ value on the test set for the full model and the full model with a single component removed for each of the ten cross-validation folds. This produced a distribution of ten R^2^ values for the full model and ten R^2^ values for the full model without a particular component. We then used a paired, one-sided Wilcoxon signed-rank test to determine if the full model explained significantly more variance than the full model without a particular component (*p < .05*). Additional criteria were applied to the intrinsic component to ensure that it was well fit by an exponential function as explained in the subsequent section.

### Estimating intrinsic and seasonal timescales

To estimate intrinsic and seasonal timescales for each neuron, we first compute the autocorrelation function associated with the autoregressive component and then fit the autocorrelation function with an exponential.

For an autoregressive process of order p (AR(p)), the Yule-Walker equations relate the autoregressive coefficients, *a_1_, a_2_, …, a_p_,* to the autocorrelation function at lag *k* (*p(k))* as follows:

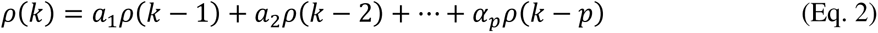

Thus, for the seasonal component, an autoregressive process of order one, the autocorrelation function at lag *k* is computed analytically, *p(k) = (a_1_)^k^* . For the intrinsic component, an autoregressive process of order ten, the Yule-Walker equations are solved numerically to obtain the autocorrelation function (using solve function in MATLAB R2022a) (**Fig. 2-2**).

An autoregressive model of order one, AR(1), has an autocorrelation function defined by a single exponentially decaying process, when the coefficient is positive. An AR(1) model with a negative coefficient has an autocorrelation function that oscillates with an exponentially decaying magnitude. For seasonal components with a negative coefficient, the timescale corresponds to the timescale of exponential decay of the magnitude of the oscillation. Thus, regardless of the sign of the seasonal coefficient, the seasonal timescale for a single neuron, *r_seasonal_*, can be defined as follows,

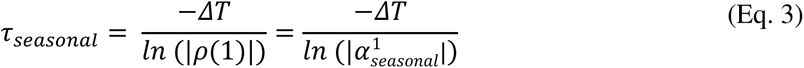

where *iJT* is the time lag between trials (here, *iJT* = 5 sec). Most coefficients of the seasonal component were positive before (188/205 = 91%) and after (236/245= 96%) training.

Nonetheless, we included neurons with both positive and negative seasonal coefficients because they both capture an exponential decay.

Because the intrinsic component is composed of ten autoregressive coefficients, it is defined by multiple exponentially decaying processes. To obtain a single intrinsic timescale, *r_intrinsic_*, we fit an exponential decay function to the autocorrelation function for each neuron:

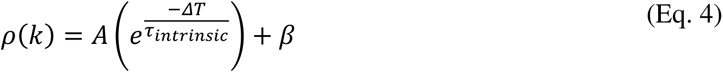

where *A* is the amplitude (fitted), *f]* is the offset, and *iJT* is the bin size (here, *iJT = 50 ms*). To mitigate the influence of refractoriness on fitted timescales, fitting started at the first reduction in autocorrelation (Wasmuht et al., 2018).

For analyses of intrinsic timescales, we selected criteria similar to Wasmuht et al. (2018) to exclude neurons whose autocorrelation function was poorly fit by an exponential function. We excluded neurons if: (1) The autocorrelation function did not decrease within 150 ms; (2) The exponential function fit had R^2^ < 0.5; or (3) The autocorrelation function did not decay towards 0 within 500 ms (*p(10) > 0.3)*. Out of 995 pre-training neurons and 904 post-training neurons that were initially classified as neurons with an intrinsic component, 954/995 and 878/904 met criterion one, 827/995 and 792/945 met criteria one and two, and 820/995 and 786/904 met all three criteria. Because these criteria resulted in the exclusion of a small proportion of neurons with the intrinsic component (<20%), all intrinsic timescale analyses were consistent with and without applying these criteria. Therefore, all neurons that met these criteria were included even if some of the coefficients in their intrinsic component were negative.

### Decoders

To determine whether intrinsic and seasonal timescales affect encoding of cue location, we trained max correlation coefficient decoders to predict cue location (9 possible locations) using firing rates of pseudo populations of neurons (see more details below). For each neuron, we randomly selected 10 trials from each cue condition (90 trials total) to use for decoding. For decoders used to study the effect of population size on decoding accuracy (**Fig. 9**), we used the average firing rate across the cue delay period as a predictor. For decoders used to analyze decoding accuracy over time (**Fig. 8**), we used the average firing rate in 500-ms sliding windows at 50-ms intervals across the fixation, cue stimulus, and cue delay periods as predictors.

We split the set of pre-training (post-training) neurons that included intrinsic timescales into two groups based on whether the neuron’s intrinsic timescale was above (long intrinsic) or below (short intrinsic) the median intrinsic timescale for the combined pre- and post-training timescale distribution (median intrinsic timescale = 91.4 msec). We followed the same procedure for neurons that included seasonal timescales (median seasonal timescale = 2.28 sec).

We trained and tested a max correlation classifier (Meyers et al. 2012) using stratified ten-fold cross-validation such that 81 trials were used in the training set and 9 trials in the test set, one for each cue condition. To train the classifier, the firing rates in the 9 trials in the training set corresponding to each cue location were averaged to construct a template vector for each cue location. Trials in the test set were then classified based on the maximum Pearson correlation between each template and the firing rate in that trial. We generated pseudo-populations of neurons for decoding by randomly sampling n neurons from neurons with long and short intrinsic and seasonal timescales (n=1-90 for **Fig. 9**, n=50 for **Fig. 8**). We repeated this resampling process 200 times to obtain a distribution of test set decoding accuracies across different pseudo-populations. To determine whether decoding accuracy was above chance, we repeated this process with shuffled cue locations. For standard error calculations, standard deviation was computed across the 50 pseudo-populations, and sample size was taken to be the number of independent pseudo-populations in the total population (e.g., 395 long intrinsic neurons/50 neurons per pseudo-population = 7.9 independent pseudo-populations).

To estimate the dimensionality of encoding, we fit an exponential function to the relationship between population size and decoding accuracy. We defined the dimensionality of the population as the minimum number of neurons where test accuracy is within 95% of its maximum. The maximum test accuracy was defined as the fitted bias parameter in the exponential function.

## Results

### Experimental paradigm and neural recordings

We obtained single-cell recordings from several subregions of the PFC (**Fig. 1a**) of monkeys before and after they learned to perform a spatial, match/nonmatch working-memory (WM) task. Before learning the main task (pre-training phase), the monkeys only had to fixate on a central point while passively viewing two stimuli separated by a delay period (delay 1 referred to as cue delay; **Fig. 1b**). After learning the main task (post-training phase), the monkeys viewed the same sequence of events in the trial but then made a saccade to one of two options to indicate whether the locations of the two presented stimuli matched or did not match (**Fig. 1c**). These features made this task particularly well-suited for studying the effects of training on fine-grained gradients of timescales in the PFC. Overall, we recorded from 2,975 neurons in the lateral PFC that we grouped into anterior-dorsal (287 neurons pre-training and 122 neurons post-training), mid-dorsal (648 and 434 neurons), posterior-dorsal (145 and 204 neurons), anterior-ventral (92 and 102 neurons), and posterior-ventral (500 and 441 neurons) PFC subregions. The data analyzed here have been previously used to examine anterior-posterior gradients of plasticity in the PFC (Riley et al., 2018).

### Model for simultaneous estimation of timescales and selectivity

To compare timescales of the dynamics of neural response in the PFC before and after training in a cognitive task, we adapted a comprehensive method to simultaneously estimate multiple timescales and selectivity from the activity of individual neurons (Spitmaan et al., 2020) during a spatial working-memory task. Similar to Spitmaan et al. (2020), the model predicts the firing rate of individual neurons in a 50-ms epoch based on activity in earlier epochs of the same trial, activity from the same epoch of previous trials, and exogenous signals (e.g., cue location) in the current trial (**Fig. 2**). This enables simultaneous estimation of intrinsic and seasonal timescales along with selectivity to exogenous signals.

The dependence of activity on recent fluctuations in neural response (i.e., intrinsic dynamics) was captured with an autoregressive (AR) component that used a weighted average of spike counts in the ten previous 50-ms epochs to predict spike count in the current epoch (AR component of order 10). Seasonal dependencies were included with an autoregressive component that used the spike count in the same epoch of the previous trial to predict spike count in the current epoch of the current trial (AR component of order 1) (**Fig. 2b**). The orders of AR components were determined based on partial autocorrelation functions (**Fig. 2-1**). To capture selectivity to exogenous signals, we incorporated components related to cue stimulus location, sample stimulus location, match vs non-match, and their interaction (**Fig. 2a**). The cue and sample components included coefficients for each of the nine possible stimulus locations. Each component had two sets of coefficients to capture selectivity separately during stimulus presentation and the delay periods (**Fig. 2a**). The coefficients from the autoregressive components were then used to estimate corresponding timescales, and the coefficients from exogenous components were used to determine selectivity for individual neurons (see Materials and Methods). To enable comparison with previous studies, we estimated intrinsic timescales by transforming autoregressive coefficients to an autocorrelation function, and then fit the resulting autocorrelation function with an exponential (**Fig. 2-2**).

**Figure 2.**
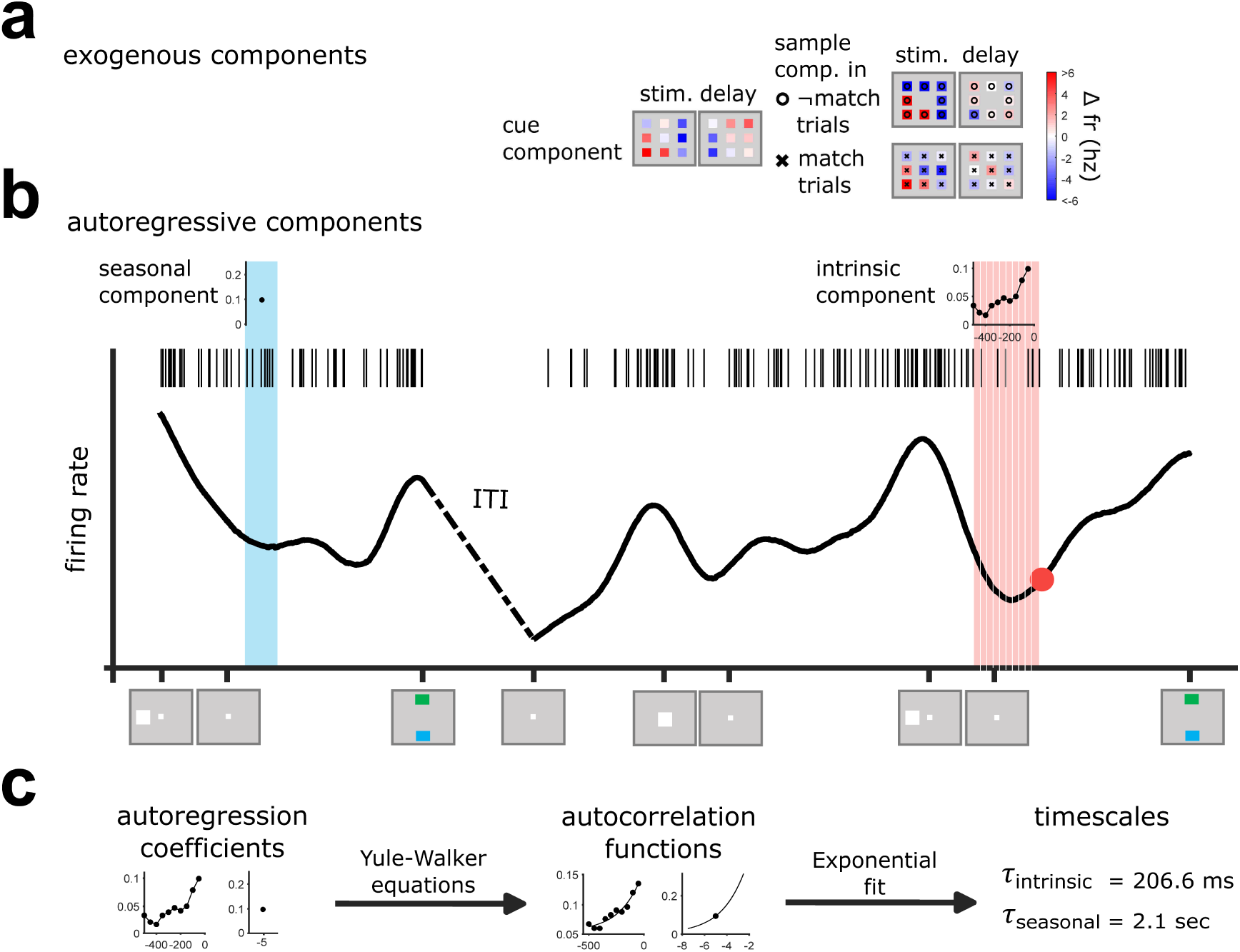
Schematic of the model for simultaneous estimation of neural timescales and selectivity in the working memory task. The model predicts the activity (number of spikes) of individual neurons in a 50-msec bin using exogenous components (a) and autoregressive components (b). **(a)** Exogenous components capture selectivity to cue, sample, match, and their interaction. The cue component captures changes in firing rate associated with each cue location and the sample+match components capture changes in firing rate associated with each sample location in match and non-match trials. **(b)** Plot shows the smoothed firing rate trace and spike raster of an example neuron across two trials. The red dot denotes the time bin for which the activity is being predicted. Activity in a given bin is dependent on activity in previous epochs in the same trial (intrinsic component, light red region) and activity in the same epoch in the previous trial (seasonal component, light blue region). Insets at the top show coefficients for the intrinsic and seasonal autoregressive components for the example neuron. **(c)** Schematic of timescale estimation. Autoregressive coefficients are mapped to autocorrelation functions (ACFs) using the Yule-Walker equations, and the ACFs are then fit with an exponential to estimate timescales (see Materials and Methods). The partial autocorrelation functions of intrinsic and seasonal dependencies can be found in Extended Data Figure 2-1. The autocorrelation functions associated with intrinsic and seasonal components can be found in Extended Data Figure 2-2.

### Anterior-posterior gradients of timescales in the response of PFC neurons during the working-memory task

Although a number of studies have established a hierarchy of intrinsic and seasonal timescales across cortical regions (Bernacchia et al., 2011; Murray et al., 2014; Spitmaan et al., 2020; Song et al., 2023), it is unclear whether more fine-grained gradients exist within individual cortical regions. To address this question, we examined the distributions of intrinsic and seasonal timescales across the anterior-posterior axis of lateral PFC neurons recorded during the spatial WM task.

To determine whether there are gradients of intrinsic and/or seasonal timescales along the anterior-posterior axis of the PFC, we fit a general linear model (GLM) to predict the estimated timescales of all neurons in dorsal (ventral) PFC using their PFC subregion (anterior = 1, middle = 2, posterior = 3) as a predictor. If the slope of this regression is significant, it indicates that timescales increase or decrease along the anterior-posterior axis of dorsal (ventral) PFC.

We found that after training in the WM task, intrinsic timescales systematically decreased along the anterior-posterior axis of the PFC in both dorsal and ventral subregions. The median value of intrinsic timescales decreased from ∼117 msec in posterior-dorsal PFC to ∼58 msec in anterior-dorsal PFC and from ∼105 msec in posterior-ventral PFC to ∼37 msec in anterior-ventral PFC (GLM for dorsal subregions: *b = −38.78, t(461) = −4.16, p = 3.79e − 5;* GLM for ventral subregions: *b = −54.57, t(325) = −3.88, p = 1.26e − 4*; **Fig. 3a**). In contrast, seasonal timescales systematically increased along the anterior-posterior axis. The median value of seasonal timescales increased from ∼2.2 sec in posterior-dorsal PFC to ∼2.4 sec in anterior-dorsal PFC and from ∼2.2 sec in posterior-ventral PFC to ∼2.4 sec in anterior-ventral PFC (GLM for dorsal subregions: *b = 0.15, t(134) = 2.33, p = 0.021;* GLM for ventral subregions: *b = 0.29, t(111) = 2.65, p = 9.15e − 3*; **Fig. 3b**). This dichotomy between intrinsic and seasonal gradients suggests that intrinsic and seasonal timescales within the PFC may arise from distinct underlying mechanisms.

**Figure 3.**
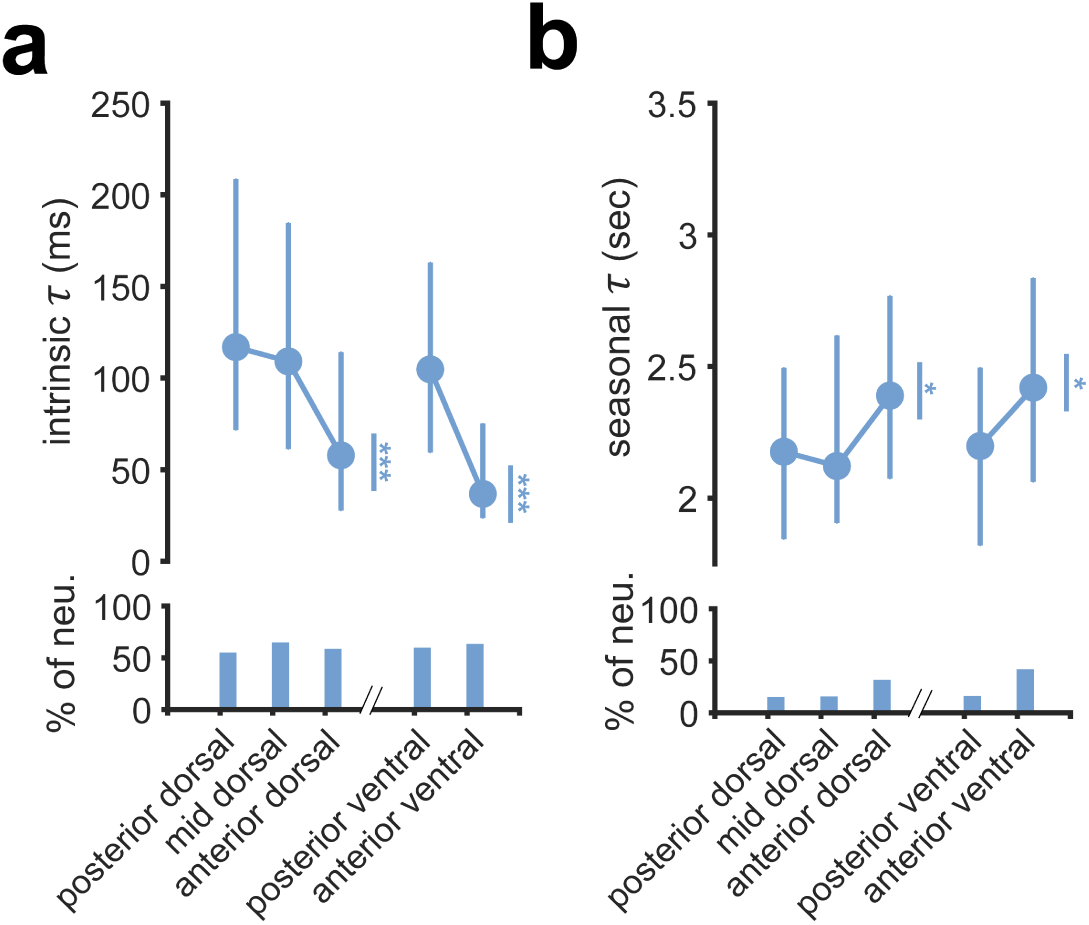
Gradients of multiple timescales across the prefrontal subregions after training in the WM task. Plots show the median of intrinsic (a) and seasonal (b) timescales in five subregions of the prefrontal cortex. Timescales were estimated using the full model for fitting the response of each neuron. Error bars indicate first and third quartiles. Insets show the fraction of neurons with a significant timescale in each subregion. Asterisks next to vertical lines indicate the significant regression slope for predicting timescales across posterior-, mid-, and anterior-dorsal subregions, or posterior- and anterior-ventral subregions (t-test of slope for regression line). In all plots, asterisks correspond to * = p<0.05, ** = p<0.005, ***=p<0.0005.

Taken together, these results demonstrate that after training in a WM task, fine-grained gradients of intrinsic and seasonal timescales exist along the anterior-posterior axis of the PFC. Although seasonal timescales parallel known anatomical and functional hierarchies of the PFC (Badre and D’Esposito, 2007, 2009; Riley et al., 2017, 2018), intrinsic timescales change in the opposite direction.

### Anterior-posterior gradient of intrinsic timescales during a passive-viewing task before training

To examine the influence of training on fine-grained gradients of timescales, we next examined timescales of neural response recorded during the pre-training task, before the monkeys learned to perform the working-memory task and instead, passively viewed cue and sample stimuli on each trial of the experiment.

We found that similar to the post-training sessions, intrinsic timescales systematically decreased along the anterior-posterior axis of dorsal PFC. The median value of intrinsic timescales decreased from ∼105 msec in posterior-dorsal PFC to ∼89 msec in anterior-dorsal PFC (GLM: b=-17.36, t(551)=-2.21, p=0.028), however, there was no evidence for a decrease from posterior-ventral to anterior-ventral PFC (GLM: b=-10.40, t(269)=-0.80, p=0.42) (**Fig. 4a**). Moreover, we did not find any evidence for a change in seasonal timescales along the anterior-posterior axis in either dorsal or ventral PFC (GLM for dorsal subregions: b=-0.09, t(147)=-1.12,p=0.26; GLM for ventral subregions: b=0.03, t(58)=0.15, p=0.88) (**Fig. 4b**).

**Figure 4.**
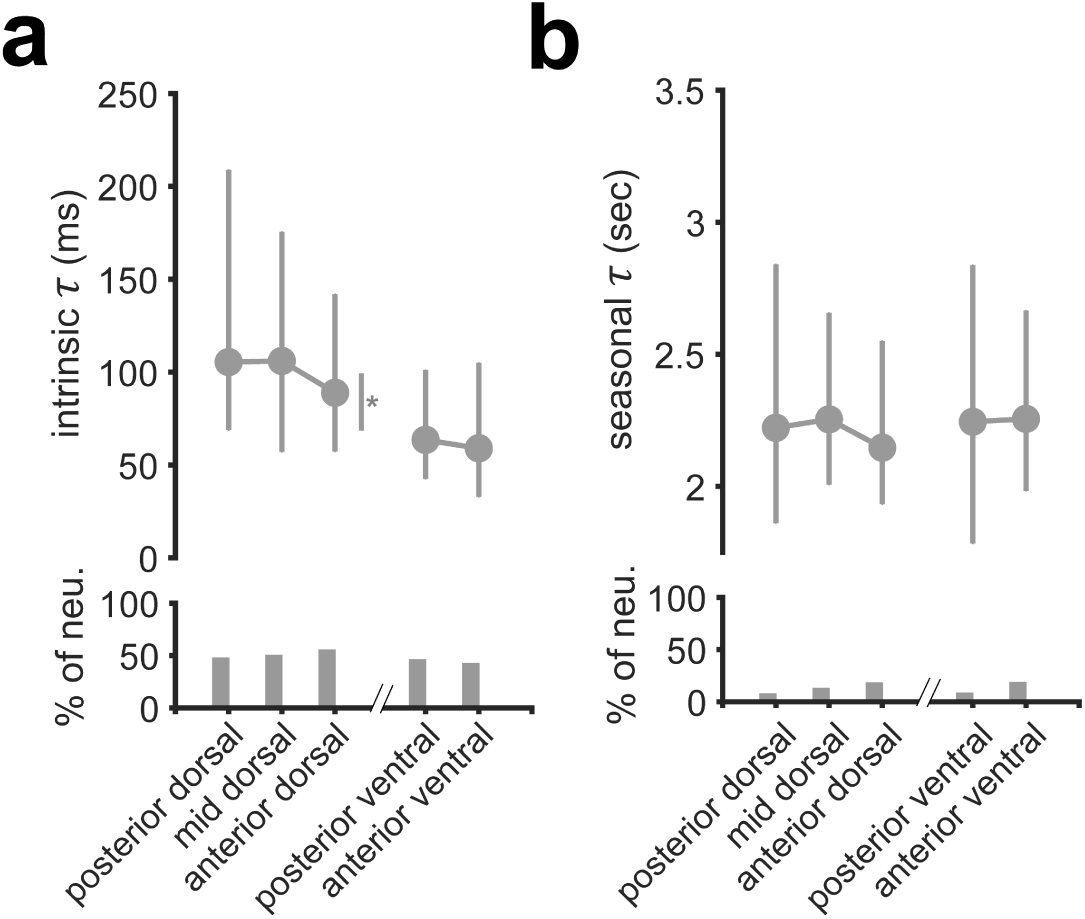
Gradients of multiple timescales across the prefrontal subregions before training in the WM task. Plots show the median of estimated intrinsic (a) and seasonal (b) timescales in five subregions of the prefrontal cortex using neural activity during pre-training sessions. Conventions are the same as in Figure 3.

Comparison of timescales across the pre- and post-training neural data indicates that training in the working-memory task could have refined the weak anterior-posterior gradient of intrinsic timescales that existed before training (but was only significant in the dorsal PFC) (**Fig. 5a**). This refinement was particularly reflected in a decrease in intrinsic timescales in anterior-dorsal and anterior-ventral PFC, and an increase in posterior-ventral (and to some extent posterior-dorsal) PFC after training (**Fig. 5a**). In addition, training was accompanied by an increase in seasonal timescales in anterior-dorsal and anterior-ventral PFC (**Fig. 5b**), resulting in gradients of timescales along the posterior-anterior axis.

**Figure 5.**
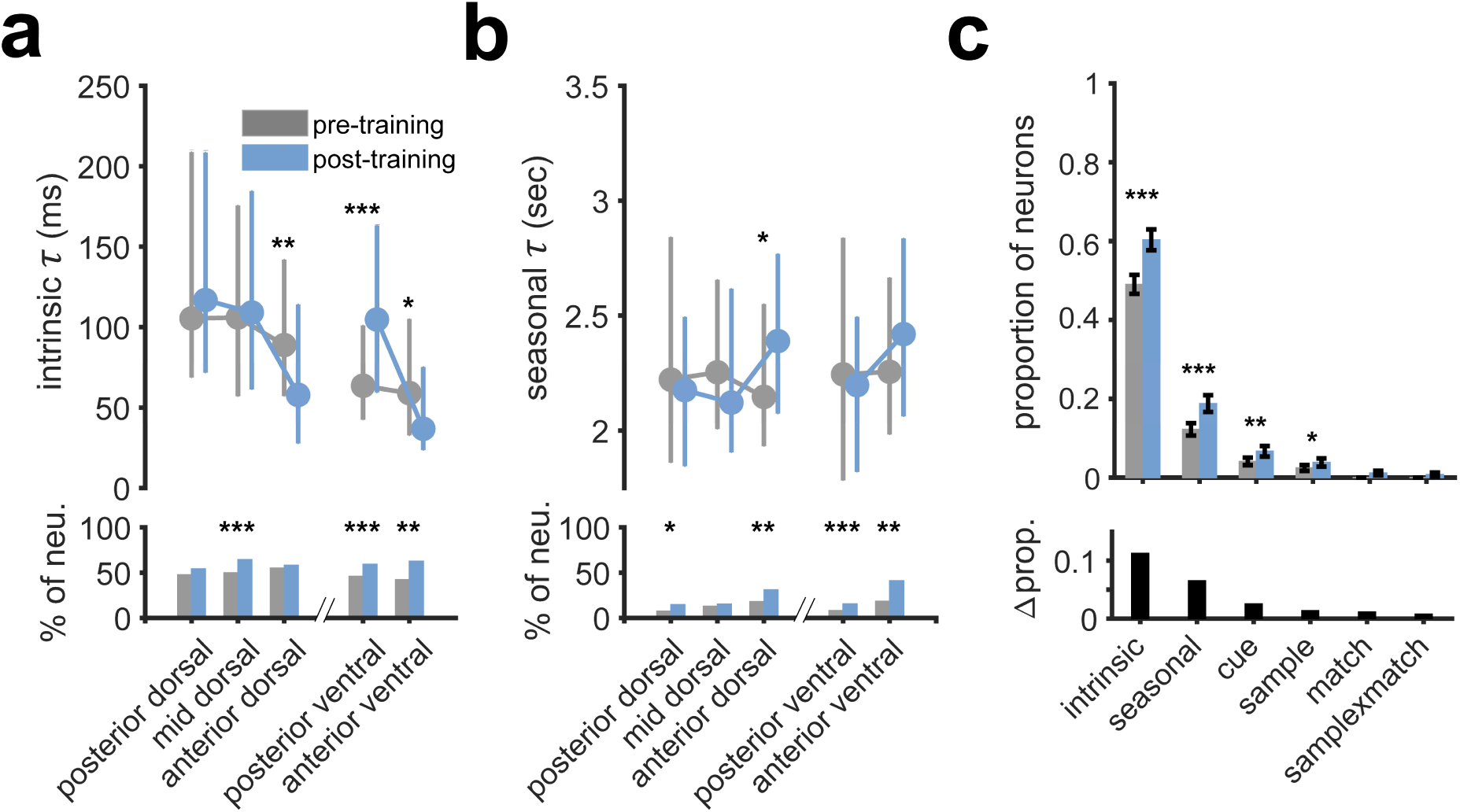
Effects of training on the magnitude and expression of timescales and on selectivity to task-relevant signals in different PFC subregions. (**a–b**) Replotted median intrinsic (a) and seasonal (b), timescales and fractions of neurons with certain timescales before (pre-training sessions; Fig. 4) and after training in the WM task (post-training sessions; Fig. 3). Asterisks on top of plots mark the significant difference between the median of timescales within a subregion between pre-training and post-training (two-sided Wilcoxon ranksum test). (**c**) Proportion of neurons with significant intrinsic, seasonal, or task-relevant components before and after training. Inset at the bottom displays the difference between the post-training and pre-training proportions. Asterisks denote a significant difference in proportion between pre-training and post-training (chi-squared test). In all plots, asterisks correspond to * = p<0.05, ** = p<0.005, ***=p<0.0005. The change in cross-validated R^2^ associated with removing each model component from the full model can be found in Extended Data Figure 5-1.

To determine whether training was associated with changes in the median or variance of timescales across the entire PFC, we compared the distribution of pre-training and post-training timescales pooled across subregions. We did not observe significant differences between the median intrinsic timescales before and after training or median seasonal timescales before and after training (two-sided Wilcoxon rank sum test; intrinsic, n = 1606, p = 0.055; seasonal, n = 450, p = 0.986). Moreover, we found that the variance of intrinsic timescales was significantly higher after training than before training (two sample F-test; n = 1606, p = 2.7e-3) whereas the variance of seasonal timescales was not significantly different before and after training (two sample F-test; n = 450, p = 0.45).

### Changes in proportions of neurons with intrinsic and seasonal timescales and with task selectivity after training

We next compared the proportions of neurons that expressed each autoregressive and task-relevant component and the variance explained by these components in pre- and post-training recordings. This analysis could reveal if training in a cognitive task caused certain dynamics to become more important for predicting neural response.

We found the proportion of neurons with all components increased after training, but these increases were significant for the proportions of neurons with the intrinsic component, seasonal component, and cue and sample exogenous components (chi-squared test; intrinsic: p=9.12e-10, seasonal: p=7.78e-7, cue: p=1.96e-3, sample: p=0.029) (**Fig. 5c**). The percentages of neurons with intrinsic timescales increased from 49% to 60% and the proportion with seasonal timescales increased from 12% to 19%. In contrast, the percentages of neurons with exogenous components increased by smaller amounts, from 4% to 7% for the cue exogenous component and from 2% to 4% for the sample exogenous component (**Fig. 5c**).

Next, to examine the importance of each model component for explaining neural activity, we computed the change in cross-validated variance associated with removing each component from the full model. We found that the full model explained ∼6% of variance in neural response before and after training in neurons that included at least one model component (pre-training mean cvR^2^ = 0.063, post-training mean cvR^2^ = 0.062), and that the removal of the intrinsic component resulted in the largest decrease in explained variance (∼3%), followed by the exogenous components, and the seasonal component (**Fig. 5-1**).

Overall, these results suggest that training was associated with increased proportions of neurons with intrinsic and seasonal neural dynamics, in conjunction with increased proportions of neurons with selectivity to task-relevant signals as reported previously (Riley et al., 2018). Among neurons with intrinsic and seasonal neural dynamics and/or selectivity to task-relevant signals, the average variance explained by these model components was similar before and after training.

### Dependence of timescales on selectivity to task-relevant signals

To determine whether estimated timescales depend on the selectivity to task-relevant signals, we next compared the distribution of intrinsic and seasonal timescales between neurons with and without selectivity to each task-relevant signal. To capture selectivity to task-relevant signals, our models included four exogenous components (for cue location, sample location, match status, and sample location × match status on the current trial). Through model selection, we identified whether the addition of an exogenous component resulted in a significant increase in explained variance. For example, if the addition of the cue component to a model containing all other components resulted in a significant increase in cross-validated explained variance, that neuron was classified as a neuron with the cue component (includes component). Otherwise, the neuron was classified as not including the component (no component). This criterion was selected to minimize false positives; i.e., neurons classified as including a component that they do not actually have, and thus some of the neurons classified as ‘not including’ a task-relevant component may still exhibit weak selectivity. Critically, because of concurrent estimation of selectivity and timescales, our method avoids mixing these two factors. In other words, our approach distinguishes variance in neural response due to the task-relevant signals in the current trial from variance in neural response due to integration of signals over time.

We found that intrinsic timescales were longer in neurons that were selective to either cue location or sample location, in both pre-training and post-training recordings (Wilcoxon ranksum test; pre-training; cue: p=4.9e-3, sample p=0.01; post-training; cue: p=2.25e-4, sample: p=4.5e-3) (**Fig. 6a,c**). In pre-training recordings, the median intrinsic timescale was lower in neurons without the cue component than those with the cue component (∼84 ms vs. ∼111 ms) (**Fig. 6a**). Similarly, in post-training recordings, the median intrinsic timescale was lower) in neurons without the cue component compared to those with the cue component (∼95 ms vs. ∼136 ms) (**Fig. 6c**). In addition, neurons with selectivity to match or sample×match exhibited longer intrinsic timescales that those without such selectivity but this effect was only significant for the former (Wilcoxon rank-sum test; post-training; match: p=0.019, sample×match p=0.053).

**Figure 6.**
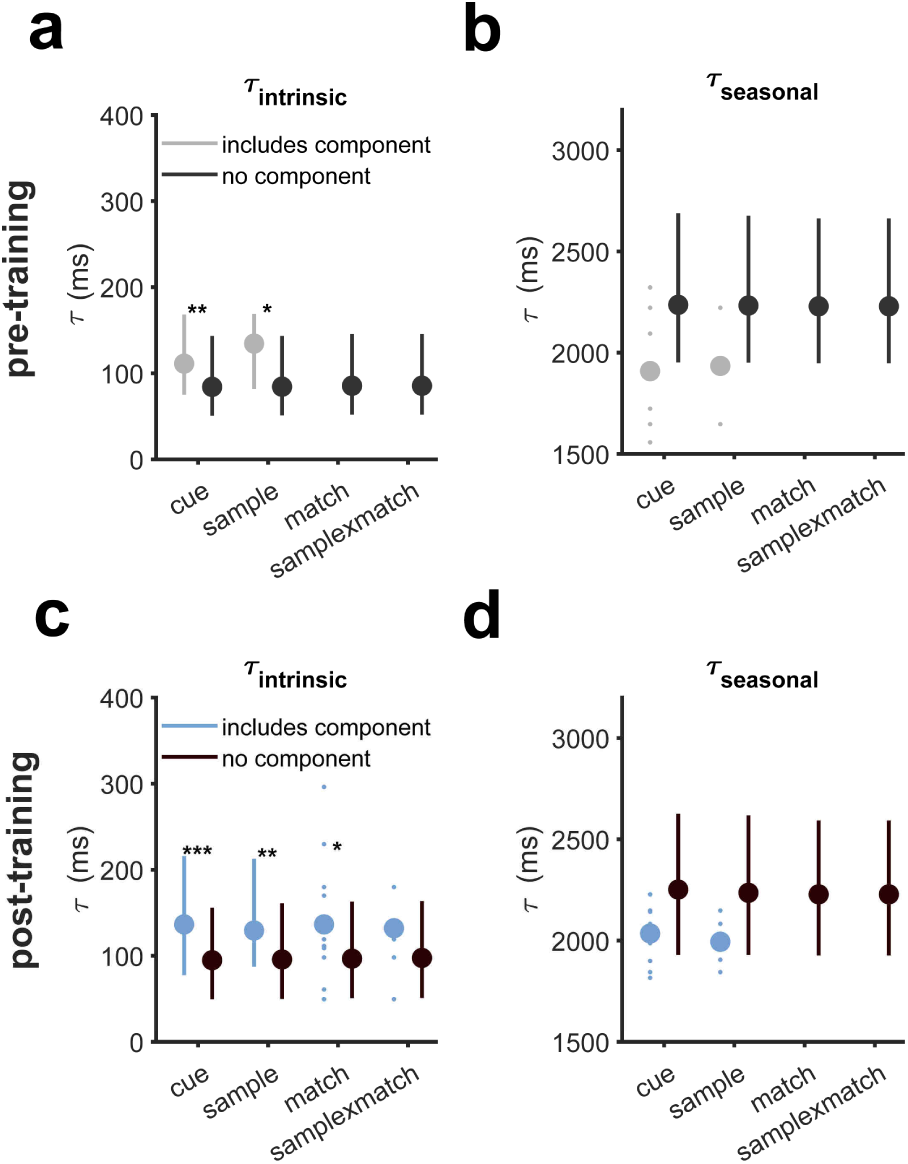
Neurons with task-relevant components have longer intrinsic timescales and a trend towards shorter seasonal timescales. Plotted is the median intrinsic (a,c) and seasonal (b,d) timescales for neurons that have a particular task-relevant component (include component) versus neurons that do not have that task-relevant component (no component) before training (a,b) and after training (c,d) Error bars denote the first and third quartiles, and individual data points are shown for conditions with fewer than 25 neurons. Asterisks mark a significant difference between the median of timescales between include component and no component conditions (two-sided Wilcoxon ranksum test; * = p<0.05, ** = p<0.005, ***=p<0.0005).

In contrast, median seasonal timescales tended to be shorter in neurons selective to cue location and sample location in both pre-training and post-training recordings, however, these differences were not statistically significant due to the small number of neurons that had seasonal timescales and included task-relevant components (Wilcoxon rank-sum test; pre-training; cue: p=0.07, sample p=0.30; post-training; cue: p=0.07, sample: p=0.17) (**Fig. 6b,d**).

Moreover, to examine the relationship between estimated timescales and performance in the task, we computed the correlation between timescales and the per-session error rate for each post-training neuron. However, we did not find evidence for significant correlations between the error rate and intrinsic or seasonal timescales (Spearman correlation; intrinsic, r = 0.03, p = 0.41; seasonal, r = -0.04, p = 0.57).

### Relationship between intrinsic and seasonal timescales and encoding of cue and sample locations

Because on average, intrinsic timescales were longer and seasonal timescales were shorter in neurons that were selective to cue and sample location than in neurons that were not selective, we hypothesized that the length of intrinsic timescales may be associated with the strength of cue-location and sample-location encoding. To test this hypothesis, we fit models composed of only the cue component or only the sample component to each neuron to assess the variance in response explained by cue location and sample location independently. We then examined the correlation between intrinsic and seasonal timescales and the variance explained by cue and sample locations to determine whether intrinsic and seasonal timescales predict encoding strength.

We found small but significant correlations between intrinsic timescales and variance explained by cue-location and sample-location components in both pre-training and post-training recordings, such that longer intrinsic timescales predicted increased variance explained by both of these exogenous components (Spearman correlation: pre-training cue: r=0.16, p=1.8e-3, pre-training sample: r=0.14, p=9.5e-3, post-training cue: r=0.23, p=4.5e-6, post-training sample: r=0.21, p=3e-5) (**Fig. 7a–b,e–f**). In contrast, we did not observe significant correlations between seasonal timescales and variance explained by cue-location or sample-location component in either pre-training or post-training recordings (Spearman correlation: pre-training cue: r=-0.15, p=0.21, pre-training sample: r=0.01, p=0.94, post-training cue: r=0.01, p=0.88, post-training sample: r=-0.07, p=0.48) (**Fig. 7c–d,g–h**). Overall, these results suggest that intrinsic timescales may be important for processing of external information even before training on a cognitive task.

**Figure 7.**
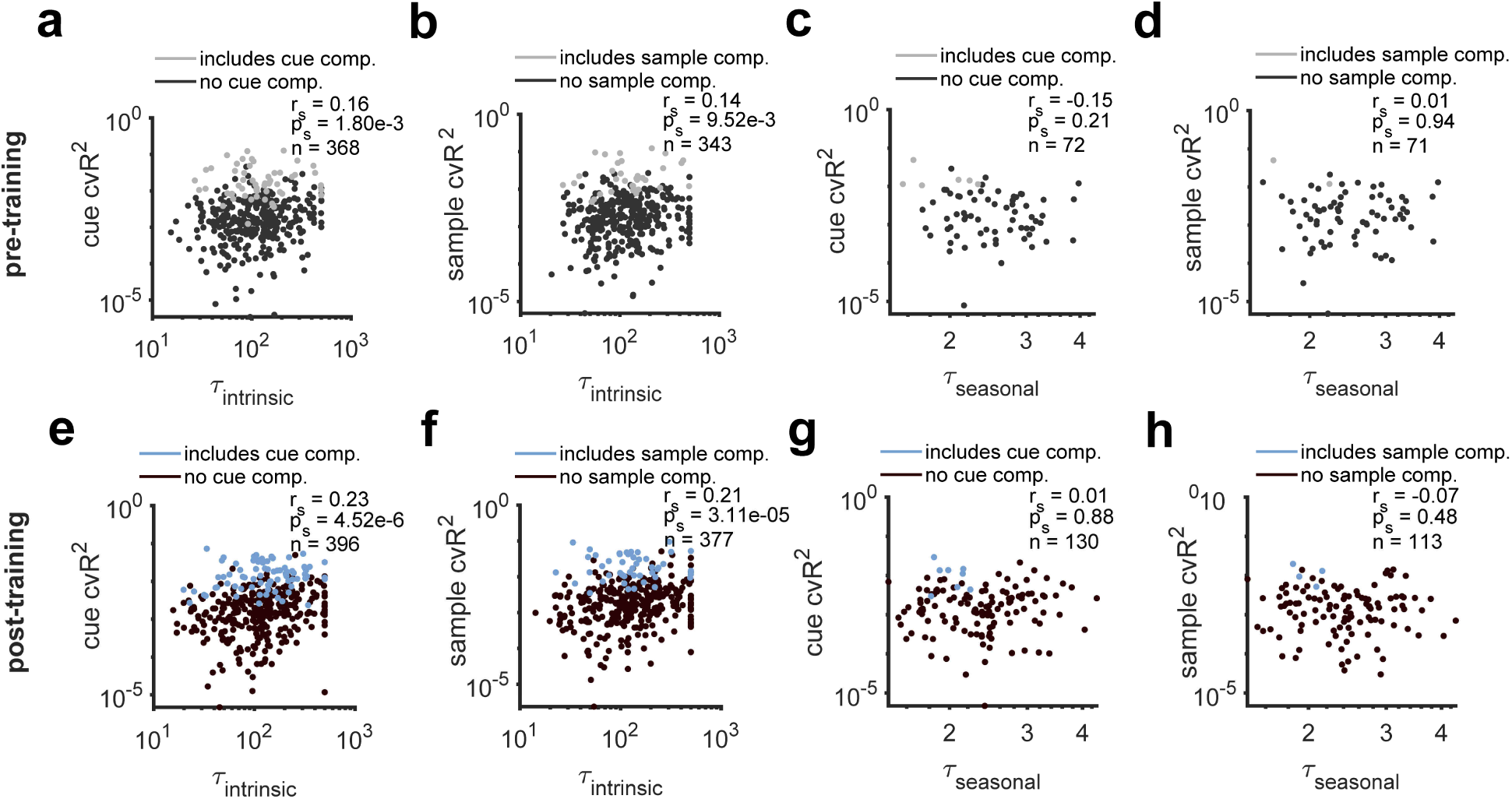
Longer intrinsic timescales are associated with more variance in response explained by cue location and sample location. Plotted is the correlation between intrinsic and seasonal timescales and variance explained by cue (a,c,e,g) and sample (b,d,f,h) location for pre-training (a,b) and post-training (c-d) neurons. Variance is defined as the cross-validated R^2^ (cvR^2^) of a model predicting neural response using only the cue location (a,c) or only the sample location (b,d). Only neurons with a significant intrinsic (seasonal) component and a cue or sample cvR^2^ greater than zero are included. Points are colored based on whether a neuron has a significant cue or sample component in the full model. The spearman correlation coefficient (r_s_) and associated p-value (p_s_) are displayed.

### Distinct contributions of intrinsic and seasonal timescales to dynamic encoding of cue location at population level

Observing the opposite dependence of intrinsic and seasonal timescales on selectivity to task-relevant signals at the single-cell level (**Fig. 6**), we next asked whether populations of neurons with short or long intrinsic timescales (similarly for seasonal timescale) encode task-relevant signals differently. To that end, we examined performance for decoding the location of the cue stimulus throughout the trial by creating pseudo-populations of 50 neurons and decoding cue-location using a max correlation classifier (see Materials and Methods for more details). We considered cue encoding because information about cue location was most relevant for task performance. Differences in decoding performance across the trial could clarify the role of timescales in encoding information during stimulus presentation versus maintaining information in working memory.

We found that the decoding performance of cue location sharply increased during stimulus presentation but mainly in populations consisting of neurons with long intrinsic timescales (**Fig. 8a–b**) or populations consisting of neurons with short seasonal timescales (**Fig. 8c–d**) both before and after training (mean +/- s.e.m. at 1250 ms; pre-training intrinsic long: 0.30 ± 0.03, short: 0.24 ± 0.02, seasonal long: 0.13 ± 0.03, short: 0.18 ± 0.04; post-training intrinsic long: 0.33 ± 0.03, short: 0.23 ± 0.03, seasonal long: short: 0.18 ± 0.03). This confirms our single-cell results suggesting that longer intrinsic timescales and shorter seasonal timescales could improve encoding of information at both single-cell and population levels.

**Figure 8.**
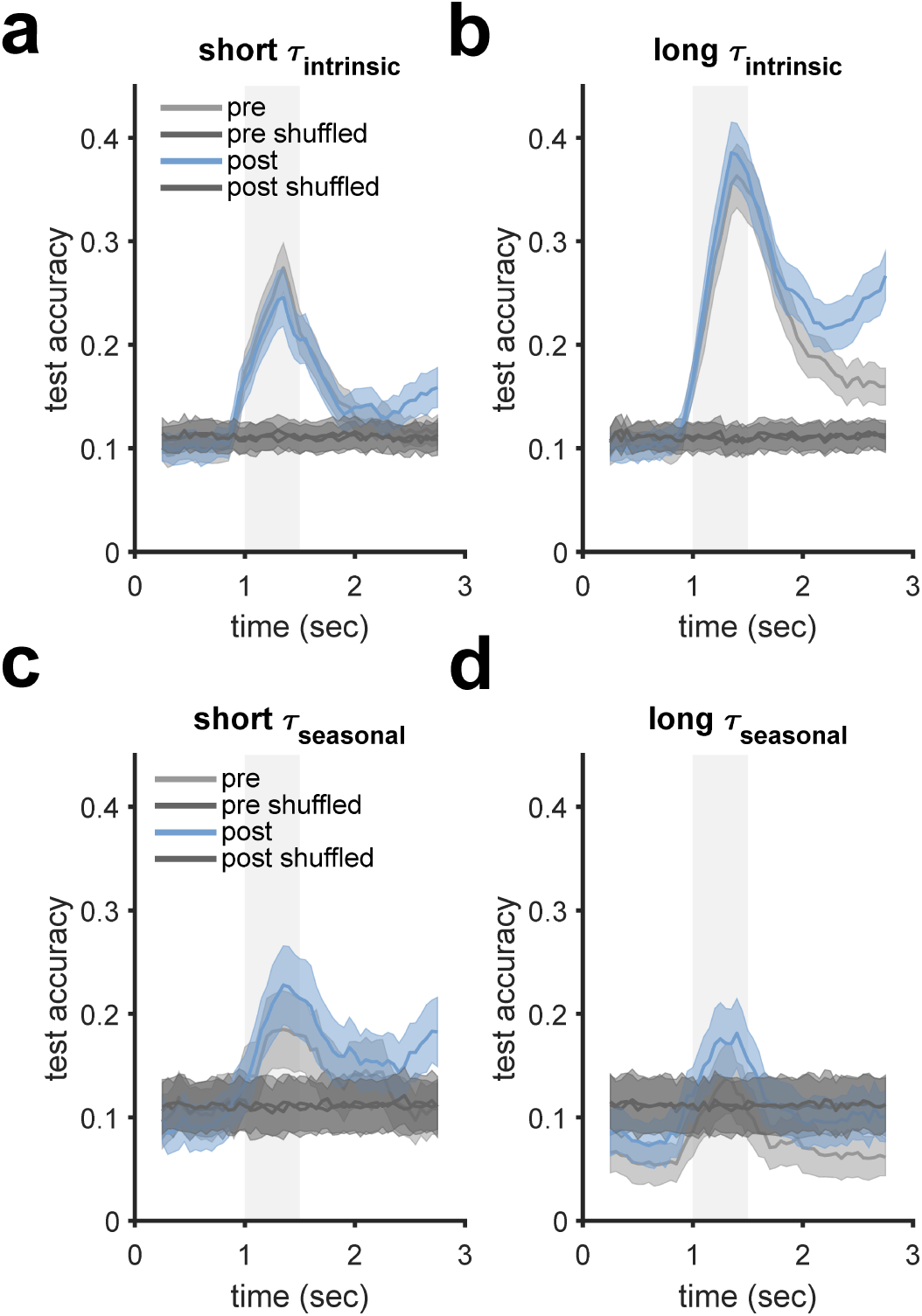
Decoding cue location over time from populations of neurons with long and short intrinsic and seasonal timescales. **(a–b)** Plotted is the test set accuracy of the cue-location decoder across time in the fixation period (0-1s), cue stimulus period (1-1.5s), and cue delay period (1.5-3s) for neurons with short (a) and long (b) intrinsic timescales. Curves denote mean decoding accuracy over 200 randomly sampled pseudo-populations of 50 neurons. Error bars denote 1 s.e.m. **(g–h)** Similar to (e–f) for short and long seasonal timescales.

More importantly, the decoding advantages of long intrinsic timescales extended to the delay period between cue and sample stimulus presentations but mainly after training, suggesting an important role for neurons with long intrinsic timescale in sustaining task-relevant information in working memory (mean +/- s.e.m. at 2750 ms for pre-training intrinsic long and post-training intrinsic long: 0.16 ± 0.02 and 0.27 ± 0.02; **Fig. 8b**). Because the overall distributions of intrinsic timescales were similar before and after training (**Fig. 5a**), this suggests that neurons with longer intrinsic timescales may be recruited differently at the population level to allow better encoding after training.

To further assess population coding during the delay period between cue and sample presentations, we also examined decoding accuracy during this period using pseudo-populations with different numbers of neurons. Dependence of encoding on the number of neurons in the population could provide information about the properties of population coding and its dependence on timescales. As expected, decoding accuracy increased with increasing numbers of neurons in the population for short intrinsic, long intrinsic, and short seasonal neurons both before and after training. However, decoding accuracy reached its asymptotic value with a smaller number of neurons for pre-training neurons compared to post-training neurons, suggesting lower dimensionality for representation of cue location before training (pre vs post for short intrinsic: 48 vs 123 neurons; pre vs post for long intrinsic: 145 vs 173 neurons; pre vs post for short seasonal: 49 vs 84 neurons; **Fig. 9a-c**). In contrast, decoding accuracy in long seasonal neurons did not increase with increasing numbers of neurons and was consistently below the shuffled control (**Fig. 9d**).

**Figure 9.**
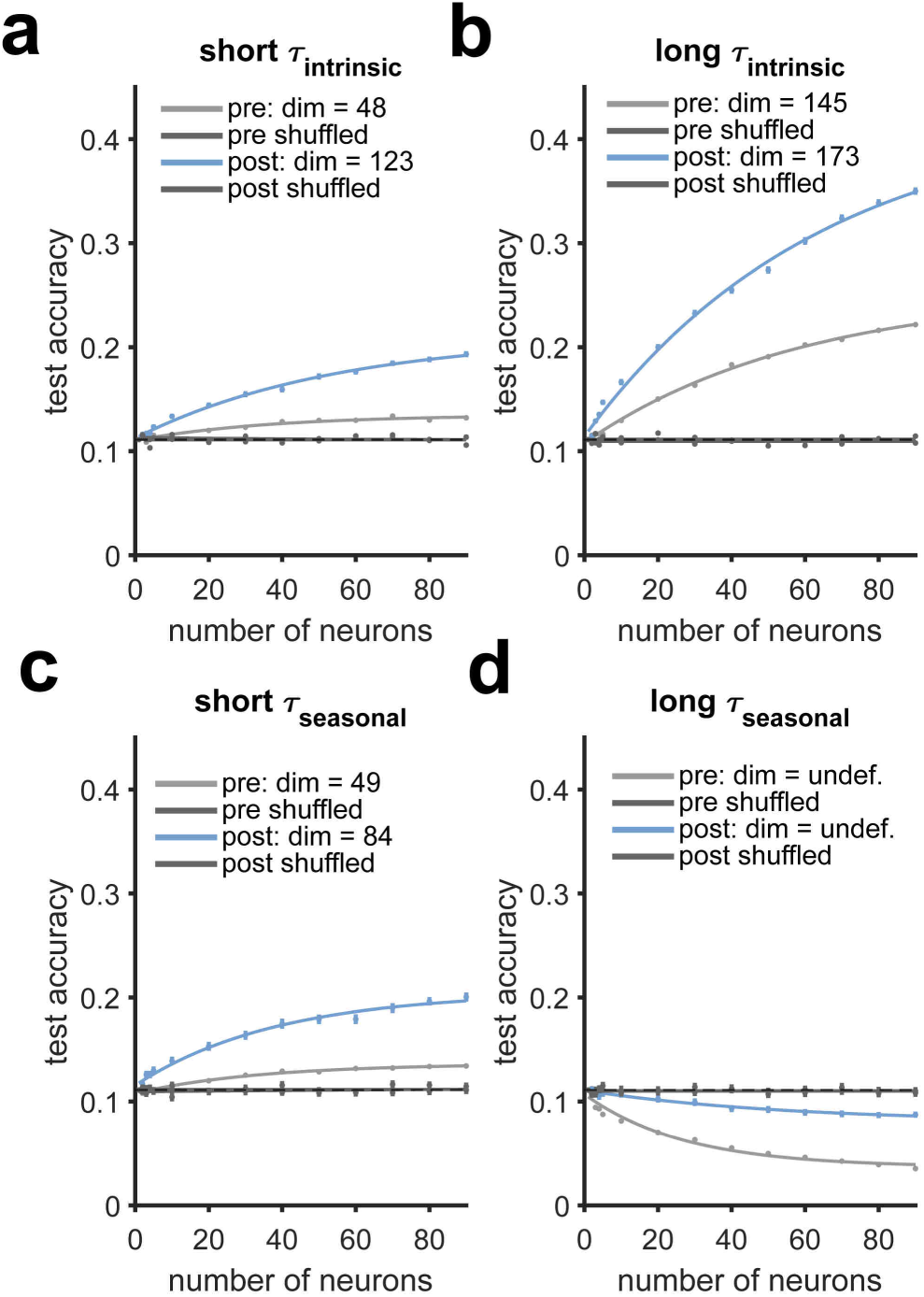
Dimensionality of cue encoding during the delay period in populations of neurons with long and short intrinsic and seasonal timescales. **(a–b)** Plotted is the test set accuracy of the cue-location decoder as a function of the size of the pseudo population used for decoding, separately for neurons with short (a) and long (b) intrinsic timescales before training (gray), after training (blue), and in shuffled controls (black). Individual points denote mean decoding accuracy over 200 randomly sampled subpopulations of a particular size. Error bars on individual points denote 1 s.e.m. Curves are exponential functions that capture the relationship between pseudo-population size and test set accuracy. The dimensionality of the population, equal to the minimum number of neurons where test accuracy is within 95% of the maximum of the exponential fit, is reported in the legend. Decoders were trained using firing rate in the cue delay period as a predictor. **(c–d)** Similar to (a–b) but for short and long seasonal timescales.

To understand why decoding accuracy in long seasonal neurons was below chance, we examined the decoder training data for individual neurons with long seasonal timescales. We found that firing rate was not stationary over the course of the session for many long seasonal neurons; for example, many of these neurons had large bursts in firing rate that lasted a few trials or increases/decreases in firing rate across the session. Because cue location was pseudo-randomized rather than fully randomized in the task, this non-stationarity coupled with weak selectivity for cue location (**Fig. 6b, d**) could lead to below chance decoding performance.

To quantify the trial-to-trial variability in response over the session, we computed the Fano factor (i.e., the ratio of variance in firing rate to mean firing rate) in neurons with different seasonal timescales during the delay period (Qi & Constantinidis 2012). We found that Fano factor was larger in long seasonal neurons compared to short seasonal neurons, and was the largest in pre-training seasonal neurons, the population that exhibited the worst decoding accuracy (two-sided t-test; pre-training long seasonal mean = 1.92; pre-training short seasonal mean = 1.40, p = 9.30e-3; post-training long seasonal mean = 1.74; post-training short seasonal mean = 1.43, p = 5.50e-3). These results are consistent with a previous study that found a decrease in Fano factor due to training (Qi and Constantinidis, 2012) and moreover, suggest a link between neural variability and seasonal timescales. Moreover, Fano factor was positively correlated with seasonal timescales (Spearman correlation; pre-training: r = 0.27, p = 1.12e-4; post-training: r = 0.31, p = 6.75e-7). Overall, we found that longer intrinsic timescales, and to some extent shorter seasonal timescales, allowed for better encoding of cue location at both single-cell and population levels, and this improvement was accompanied by selective changes in variability and representations.

## Discussion

Using a comprehensive method to simultaneously estimate timescales and selectivity from the activity of individual neurons before and after training on a WM task, we identified fine-grained gradients of timescales within the PFC that mainly appeared after training. Intrinsic timescales decreased and seasonal timescales increased from posterior to anterior PFC. In addition, populations of neurons with longer intrinsic timescales and shorter seasonal timescales exhibited superior population-level encoding of the cue stimulus location. Notably, these effects were similar during stimulus presentation before and after training, but extended to the delay period between cue and sample presentations mainly after training, suggesting that neurons with long intrinsic timescales may be selectively recruited for maintaining information in working memory.

### Inherent and training-dependent gradients of intrinsic and seasonal timescales across PFC subregions during the WM task

A consensus of studies have found that intrinsic and seasonal timescales increase in parallel across the cortical anatomical hierarchy from early visual and somatosensory areas to prefrontal and cingulate cortices (Bernacchia et al., 2011; Murray et al., 2014; Pinto et al., 2022; Spitmaan et al., 2020; Soltani et al., 2021; Zeraati et al., 2023; Song et al. 2023). By illustrating similar, but opposite, fine-grained gradients of intrinsic and seasonal timescales along the anterior-posterior axis of the PFC after training on the WM task, our results expand upon previous work in three novel ways:

First, they provide evidence for fine-grained gradients of both intrinsic and seasonal timescales within the PFC. Previous studies aiming to link intrinsic timescales and WM were not able to identify clear differences in intrinsic timescales between prefrontal subregions, such as FEF and lPFC (Wasmuht et al. 2018) or dlPFC, vlPFC, and OFC (Cavanagh et al. 2018). This could be due to two reasons. First, these studies grouped anterior and posterior regions of lateral PFC together. Second, they relied on autocorrelation to measure intrinsic timescales and this method does not take into account other response properties of a neuron such as seasonal dependencies as shown previously (Spitmaan et al, 2020).

Second, most studies of neuronal timescales have measured timescales only after extensive training in a given task (Murray et al., 2014; Cavanagh et al., 2018; Wasmuht et al., 2018; Pinto et al., 2022; Spitmaan et al., 2020; Zeraati et al., 2023). By analyzing timescales before and after training on the WM task, we observed that gradients of intrinsic timescales in ventral subregions and seasonal timescales in dorsal and ventral subregions were less clear prior to training, suggesting that such gradients of timescales could be further sharpened by training. Our observation that many PFC neurons exhibit intrinsic timescales in both pre- and post-training recordings dovetails previous studies that suggest intrinsic dynamics are inherent properties of cortical neurons (Murray et al., 2014; Spitmaan et al., 2020). Nonetheless, we observed larger proportions of neurons with intrinsic and seasonal timescales following training as well as changes in timescale gradients following training, suggesting that neuronal timescales, even the “intrinsic” timescales, may be malleable. One possibility is that training changes connectivity within a circuit, resulting in new interactions between neurons that change the dynamics of response captured by intrinsic and seasonal timescales in addition to selectivity to task-relevant signals.

Finally, we found opposite gradients of intrinsic and seasonal timescales within subregions of the PFC. Interestingly, we also found that longer intrinsic timescales and shorter seasonal timescales contribute to better encoding of the cue stimulus location (more on this below). An anterior-posterior decrease in intrinsic timescales along with an anterior-posterior increase in seasonal timescales could decorrelate mechanisms by which population coding changes in these subregions, resulting in more flexibility in encoding relevant information.

### Distinct contributions of intrinsic and seasonal timescales to encoding task-relevant information

How intrinsic and seasonal timescales relate to the encoding of task-relevant information remains unclear, but recent studies have made progress in addressing this question. After training in a WM task, Cavanagh and colleagues found that representations of spatial cues and reward cues were strongest over the delay period in populations of vlPFC neurons with long intrinsic timescales (Cavanagh et al., 2018). Similarly, Wasmuht and colleagues showed that neurons with longer intrinsic timescales in lateral PFC carry more information about task-relevant signals during the delay period (Wasmuht et al., 2018). Thus, our results confirm previous findings that populations of neurons with long intrinsic timescales exhibit better encoding of information that should be sustained over time. Our findings additionally show a similar role for populations of neurons with shorter seasonal timescales and reveal that at the single-cell level, neurons with the strongest selectivity to task-relevant signals tend to have longer intrinsic and shorter seasonal timescales, thus providing a link between response selectivity and timescales of neural dynamics.

Previous studies have suggested that neurons with long intrinsic timescales could be important for performing cognitive tasks that require accumulation of evidence or maintenance of information over time (Wasmuht et al. 2018, Cavanagh et al. 2018; Pinto et al. 2022). Our finding that selectivity and long-intrinsic timescales co-occur at the single-cell level before and after training aligns with this hypothesis and moreover, provides a possible underlying mechanism. If a neuron’s response is highly correlated with its previous activity (long intrinsic timescale) and the neuron is selective to a visual stimulus in certain parts of space (strong selectivity), then its activity during the delay period following stimulus presentation will still be selective to the visual stimulus. Together, these could allow for better maintenance of the information in working memory.

Moreover, our finding that selectivity and short-seasonal timescales co-occur at the single-cell level may also have an intuitive explanation. If a neuron’s response is highly correlated with its response in the previous trial (long seasonal timescale) despite differences in stimuli across trials, then the neuron’s response must be relatively stimulus invariant (weak selectivity). We also found that seasonal timescales were positively correlated with Fano factor. Together, these results suggest that more response variability, which can disrupt coding capacity, could result in longer memory of response to task events that could be useful for other cognitive functions such as prediction of future events.

Finally, our analysis on the dimensionality of representations revealed that training was accompanied by an increase in the dimensionality of cue stimulus representation. This suggests that training could increase the coding ability of the population of neurons in the PFC perhaps by reduction in noise correlation (Qi and Constantinidis 2012; Mazzucato et al., 2016; Dehaqani et al., 2018; Engel and Steinmetz, 2019).

### Effect of training on the encoding of task-relevant signals in populations of neurons with long intrinsic timescales

A critical finding of our study is that training changes population coding for certain types of neurons during a specific part of the task. More specifically, we found that training accompanies improved encoding of cue location during the cue delay period in populations of neurons with long intrinsic timescales, despite similar encoding before and after training during visual stimulus presentation. This suggests that neurons with long-intrinsic timescales do not inherently encode information across a delay simply because they have a long-intrinsic timescale (Wasmuht et al. 2018, Cavanagh et al. 2018). Rather, following training in the task, neurons with long intrinsic timescales could be selectively recruited for the maintenance of task relevant information across the delay period. Thus, training on a working-memory task seems to accompany changes in neural dynamics that result in improvement in encoding, during the most critical epoch of the cognitive task.

### Limitations and future studies

There are a few limitations with our modeling strategy and dataset that are relevant to our interpretation of the results. On the one hand, because the model estimates selectivity and timescales simultaneously, our method does not mix these two factors. On the other hand, if there is multicollinearity between predictors in the model or if the model components fail to faithfully capture temporal dynamics in neural response to task signals, estimation of timescales may be biased. Moreover, our use of a stringent criterion for identifying neurons that include a particular model component mitigates false positives but could bias our later analyses by excluding neurons with weak selectivity or weak autoregressive dependencies. Finally, recordings before and after training were from different sets of neurons, and task demands were different before and after training despite identical visual stimuli. Thus, the changes in timescale gradients and encoding identified here are associated with training and/or task demands but not necessarily causally related to either. It is upon future studies to track timescales in individual neurons over the course of training using chronic recordings, and to identify the causal role of neurons with different timescales in cognitive tasks (Pinto et al. 2022).

Taken together, our results demonstrate that simultaneous expression of intrinsic timescales, seasonal timescales, and stimulus selectivity is an inherent characteristic of neurons across the PFC. Nonetheless, the gradients of these timescales and their role in encoding task-relevant information may be modulated by training or task demands.

## Acknowledgments

This work is supported by the National Institutes of Health (grant R01 DA047870 to AS and R01EY017077 to CC).

## Extended Data

**Figure 2-1.**
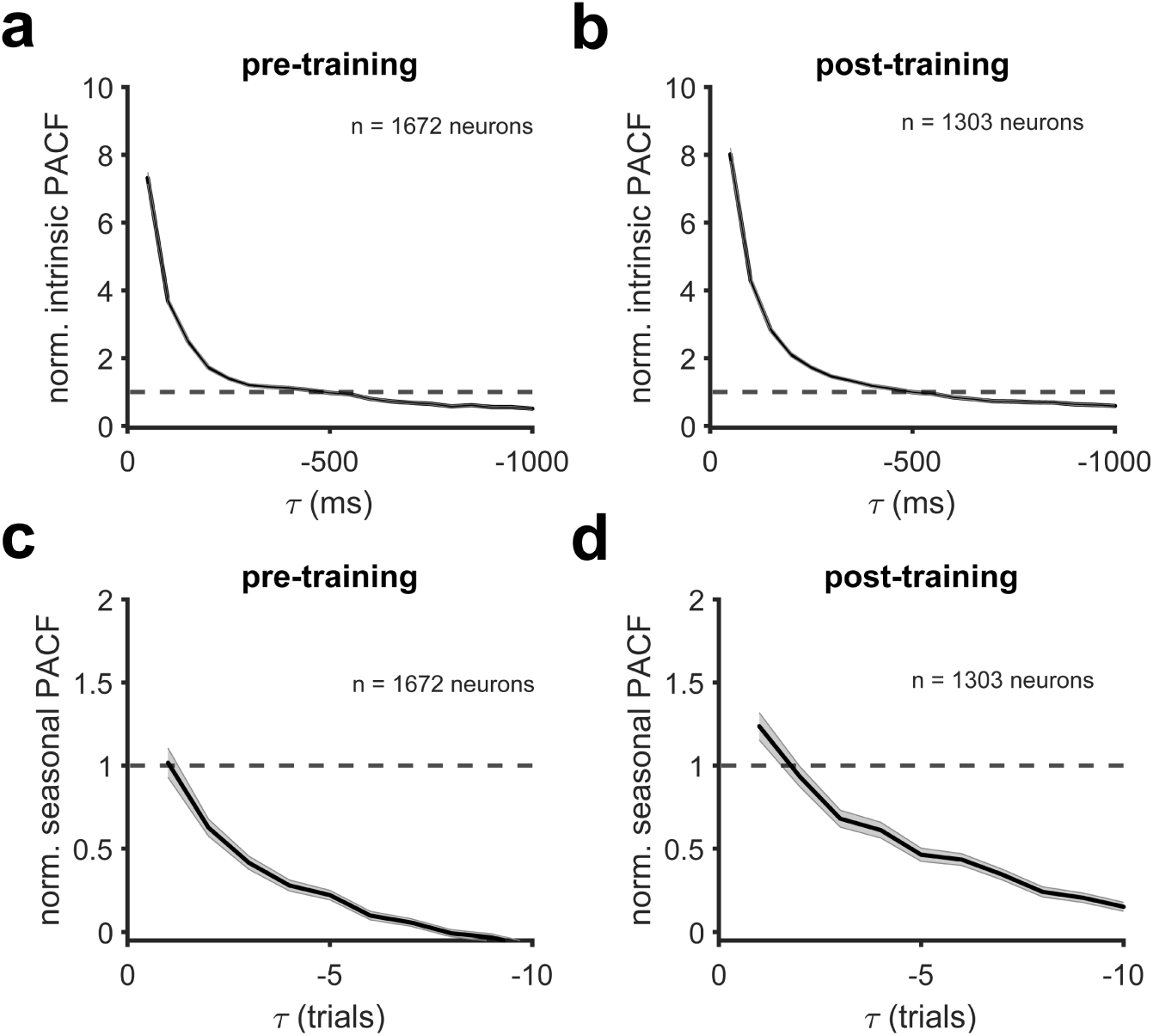
Partial autocorrelation functions of intrinsic and seasonal dependencies across all neurons. Plotted are the normalized partial autocorrelation functions (PACFs) for intrinsic (a, b) and seasonal (c, d) dependencies averaged over all neurons before training (a, c) and after training (b, d). The partial autocorrelation function for each neuron is normalized with respect to its 95% confidence interval. Dashed lines denote the upper 95% confidence bound of the PACF. Error bars denote s.e.m. The intrinsic PACFs cross the confidence bound between -500 and -550 msec, indicating that an order of 10 (10 times 50 msec bins = 500 msec) is needed for the intrinsic AR component. The seasonal PACFs cross the confidence bound between trials 1 and 2, indicating a seasonal component with order 1.

**Figure 2-2.**
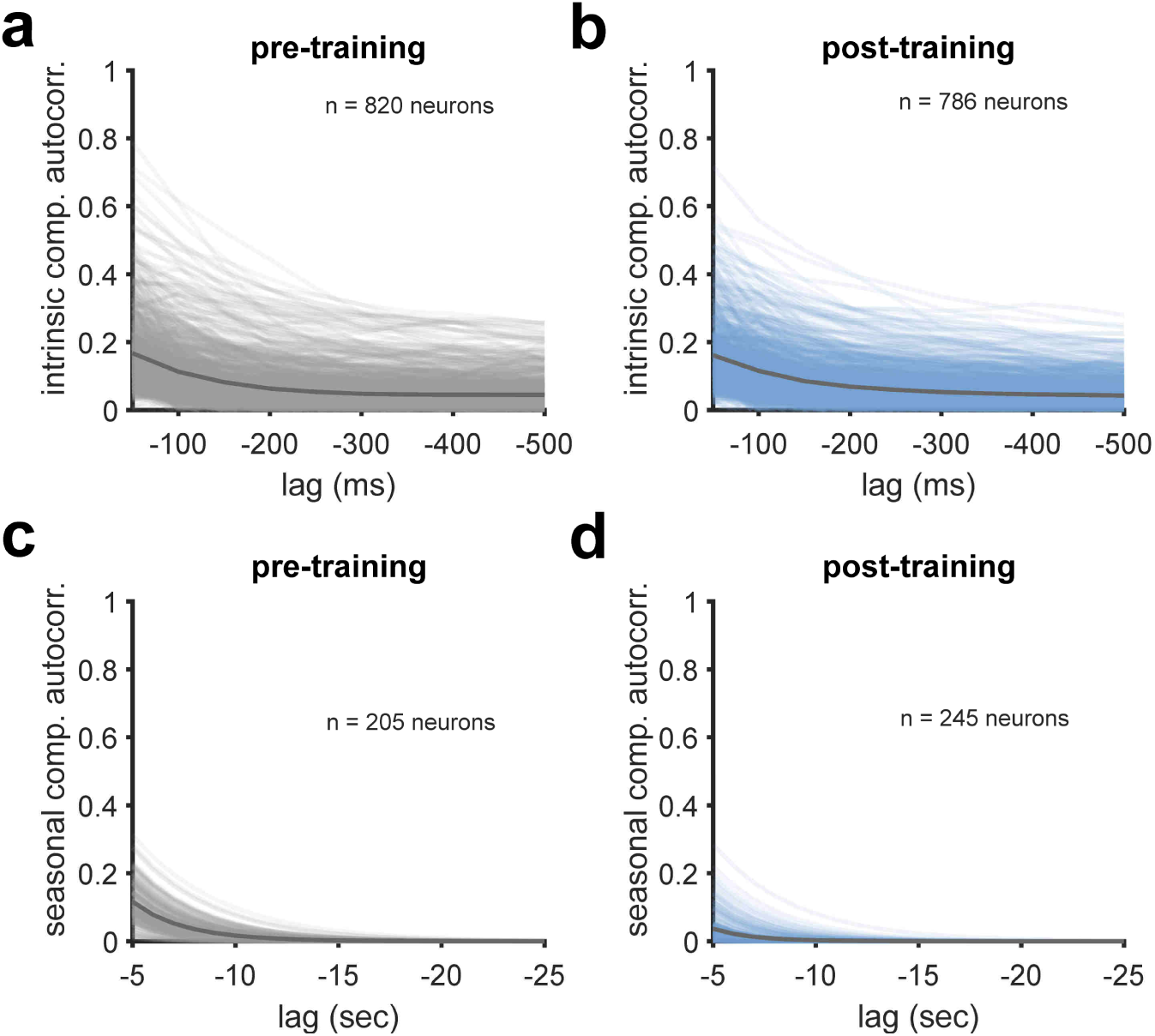
Autocorrelation functions associated with intrinsic and seasonal components. Plotted are the autocorrelation functions (ACFs) and ACF amplitudes estimated using the Yule-Walker equations from intrinsic components (a, b) and seasonal components, respectively, (c, d) before training (a, c) and after training (b, d). ACF amplitudes are plotted for the seasonal component to represent timescales of neurons with positive and negative seasonal coefficients on the same scale. Only neurons with a significant intrinsic or seasonal component are included. Dark gray curves denote the average of the autocorrelation function across all neurons. Light gray and blue curves show the autocorrelation function for individual neurons.

**Figure 5-1.**
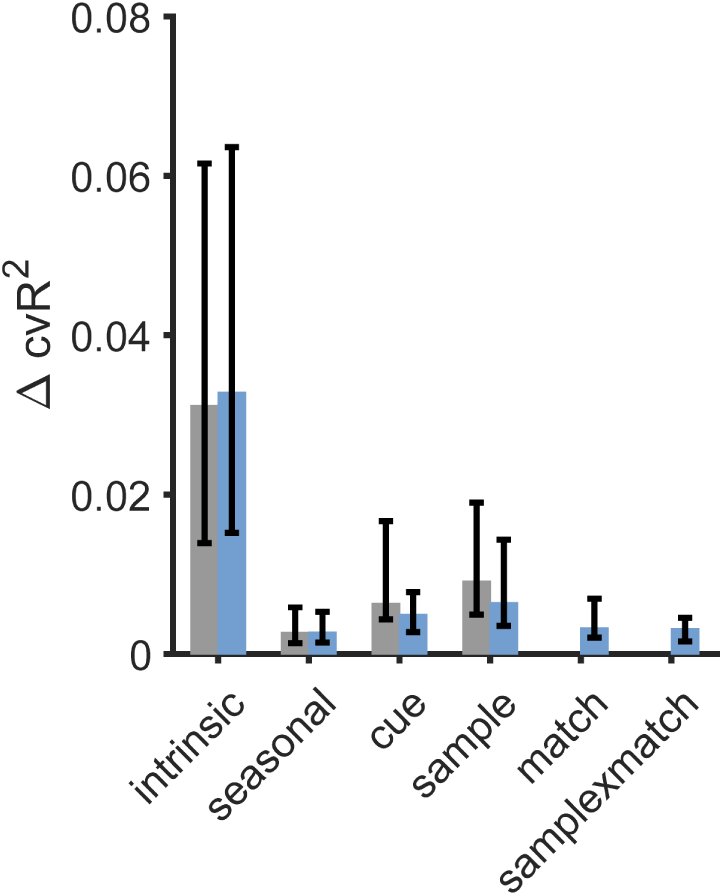
Change in cross-validated R^2^ associated with removing each model component from the full model. Plotted is the median difference in cross-validated R^2^ (cvR^2^) between the full model and the full model without a particular component in pre-training (gray) and post-training (blue) neurons. Only neurons that include a particular component are included in the plot of cvR^2^ for that particular component. Error bars denote first and third quartiles.

## References

Badre D, D’Esposito M (2007) Functional magnetic resonance imaging evidence for a hierarchical organization of the prefrontal cortex. J Cogn Neurosci 19:2082–2099.

Badre D, D’Esposito M (2009) Is the rostro-caudal axis of the frontal lobe hierarchical? Nat Rev Neurosci 10:659–669.

Bernacchia A, Seo H, Lee D, Wang X-J (2011) A reservoir of time constants for memory traces in cortical neurons. Nat Neurosci 14:366–372.

Box G, Jenkins G, Reinsel G, Ljung G (2015) Time Series Analysis: Forecasting and Control, 5th Edition. Wiley.

Brainard DH (1997) The Psychophysics Toolbox. Spat Vis 10:433–436.

Cavanagh SE, Towers JP, Wallis JD, Hunt LT, Kennerley SW (2018) Reconciling persistent and dynamic hypotheses of working memory coding in prefrontal cortex. Nat Commun 9:3498.

Chaudhuri R, Knoblauch K, Gariel M-A, Kennedy H, Wang X-J (2015) A Large-Scale Circuit Mechanism for Hierarchical Dynamical Processing in the Primate Cortex. Neuron 88:419–431.

Dang W, Jaffe RJ, Qi X-L, Constantinidis C (2021) Emergence of Nonlinear Mixed Selectivity in Prefrontal Cortex after Training. J Neurosci 41:7420–7434.

Dehaqani, MRA, Vahabie AH, Parsa M, Noudoost B, Soltani A (2018) Selective changes in noise correlations contribute to an enhanced representation of saccadic targets in prefrontal neuronal ensembles. Cerebral Cortex, 28(8), 3046–3063.

Engel TA, and Steinmetz NA (2019) New perspectives on dimensionality and variability from large-scale cortical dynamics. Current opinion in neurobiology, 58, 181–190.

Golesorkhi M, Gomez-Pilar J, Zilio F, Berberian N, Wolff A, Yagoub MCE, Northoff G (2021) The brain and its time: intrinsic neural timescales are key for input processing. Commun Biol 4:1–16.

Goris RLT, Movshon JA, Simoncelli EP (2014) Partitioning neuronal variability. Nat Neurosci 17:858–865.

Honey CJ, Thesen T, Donner TH, Silbert LJ, Carlson CE, Devinsky O, Doyle WK, Rubin N, Heeger DJ, Hasson U (2012) Slow cortical dynamics and the accumulation of information over long timescales. Neuron 76:423–434.

Kim R, Sejnowski TJ (2021) Strong inhibitory signaling underlies stable temporal dynamics and working memory in spiking neural networks. Nat Neurosci 24:129–139.

Mazzucato L, Fontanini A, La Camera G (2016) Stimuli reduce the dimensionality of cortical activity. Front Syst Neurosci. 10:11.

Meyers E, Qi X-L, Constantinidis C (2012) Incorporation of new information into prefrontal cortical activity after learning working memory tasks. PNAS 109 (12):4651–4656.

Murray JD, Bernacchia A, Freedman DJ, Romo R, Wallis JD, Cai X, Padoa-Schioppa C, Pasternak T, Seo H, Lee D, Wang X-J (2014) A hierarchy of intrinsic timescales across primate cortex. Nat Neurosci 17:1661–1663.

Pinto L, Tank DW, Brody CD (2022) Multiple timescales of sensory-evidence accumulation across the dorsal cortex. eLife 11:e70263.

Qi XL, Constantinidis C (2012) Variability of prefrontal neuronal discharges before and after training in a working memory task. PLoS One, 7(7), e41053.

Qi XL, Constantinidis C (2012) Correlated discharges in the primate prefrontal cortex before and after working memory training. Eur J Neurosci, 36(11):3538–3548.

Rigotti M, Barak O, Warden MR, Wang X-J, Daw ND, Miller EK, Fusi S (2013) The importance of mixed selectivity in complex cognitive tasks. Nature 497:585–590.

Riley MR, Qi X-L, Constantinidis C (2017) Functional specialization of areas along the anterior-posterior axis of the primate prefrontal cortex. Cereb Cortex 27:3683–3697.

Riley MR, Qi X-L, Zhou X, Constantinidis C (2018) Anterior-posterior gradient of plasticity in primate prefrontal cortex. Nat Commun 9:3790.

Seo H, Barraclough DJ, Lee D (2009) Lateral intraparietal cortex and reinforcement learning during a mixed-strategy game. J Neurosci 29:7278–7289.

Seo H, Lee D (2007) Temporal filtering of reward signals in the dorsal anterior cingulate cortex during a mixed-strategy game. J Neurosci 27:8366–8377.

Soltani A, Murray JD, Seo H, Lee D (2021) Timescales of Cognition in the Brain. Curr Opin Behav Sci 41:30–37.

Song M, Shin EJ, Seo H, Soltani A, Steinmetz NA, Lee D, Jung MW, Paik SB (2023) Hierarchical gradient of timescales in the mammalian forebrain. bioRxiv. doi:10.1101/2023.05.12.540610

Spitmaan M, Seo H, Lee D, Soltani A (2020) Multiple timescales of neural dynamics and integration of task-relevant signals across cortex. Proc Natl Acad Sci U S A 117:22522–22531.

Wasmuht DF, Spaak E, Buschman TJ, Miller EK, Stokes MG (2018) Intrinsic neuronal dynamics predict distinct functional roles during working memory. Nat Commun 9:3499.

Zeraati R, Shi Y-L, Steinmetz NA, Gieselmann MA, Thiele A, Moore T, Levina A, Engel TA (2023) Intrinsic timescales in the visual cortex change with selective attention and reflect spatial connectivity. Nat Commun 14: 1858.

